# Adenosine A2B Receptor Activation: A Novel Therapeutic Strategy for Accelerating Liver Recovery After Acetaminophen Overdose

**DOI:** 10.64898/2026.06.29.735109

**Authors:** Giselle Sanchez-Guerrero, David S. Umbaugh, Nga T. Nguyen, Hartmut Jaeschke, Anup Ramachandran

## Abstract

An acetaminophen (APAP) overdose is the leading cause of drug-induced hepatotoxicity and acute liver failure (ALF) in the United States. While N-acetylcysteine (NAC), is highly effective when administered early after an overdose, its efficacy decreases with delayed administration. Since most patients present late to the clinic, there is an urgent need for novel late-acting therapeutic options to prevent progression to ALF. We previously demonstrated the benefit of delayed activation of the Adenosine A2B Receptor (A2BAR) in attenuating APAP-induced hepatotoxicity and this study focuses on its effects on liver recovery after injury. Fasted male C57BL/6J mice were treated with 300 mg/kg APAP, followed by activation of A2BAR 6 or 9 h later and sacrifice 24, 48 or 72 h post-APAP with evaluation of liver injury, the innate immune response and liver regeneration. Delayed activation of A2BAR significantly enhanced liver recovery, with accelerated repopulation of the liver by Kupffer cells, increased macrophage migration to the necrotic areas and their faster resolution. A2BAR activation also upregulated lipid metabolism-related genes in non-parenchymal cells and cell proliferation and metabolism genes in hepatocytes. Remarkably, genes such as *Cidec* and *Plin2*, crucial for lipid droplet formation, were upregulated, indicating that A2ABR activation enhances lipid metabolism which plays a key role in providing energy for liver regeneration. Overall, these findings highlight the potential of A2BAR activation not only in protecting against liver injury, but also in promoting and accelerating liver regeneration by modulating the innate immune responses and metabolic pathways.

## 1. Introduction

Acetaminophen (APAP) is a widely used analgesic and antipyretic medication. Despite its safety and effectiveness at therapeutic doses, an APAP overdose can cause acute liver injury. This is primarily mediated by the production of a toxic metabolite, N-acetyl-p-benzoquinoneimine (NAPQI), formed in the liver by the cytochrome P450 (CYP) enzyme system. NAPQI depletes glutathione (GSH) [1], a critical cellular antioxidant, leading to oxidative stress, mitochondrial dysfunction, and ultimately, hepatocyte necrosis [2]. N-acetylcysteine (NAC), which replenishes hepatic glutathione stores, is the current standard treatment for an APAP overdose, but its effectiveness is limited by a narrow therapeutic window [3]. Although fomipezole, the new antidote under clinical development, is more effective than NAC in preventing liver injury, it uses the same therapeutic targets, i.e., NAPQI formation and oxidant stress [4, 5]. Therefore, there is an urgent need for novel therapeutic strategies to combat not only APAP-induced liver injury, but also promote regeneration, thus functioning in a delayed fashion beyond the effective window for NAC and fomepizole.

The adenosine A2B receptor (A2BAR) belongs to the family of G-protein-coupled receptors and plays a role in various physiological processes, including inflammation, cell death, and metabolism. Our recent research has shed light on the potential of A2BAR activation as an approach to protect against liver injury after an APAP overdose [6–8]. Our previous studies provide intriguing insights into the interplay between A2BAR and APAP pathophysiology. We investigated the therapeutic potential of A2BAR activation in APAP overdose using the specific A2BAR agonist, BAY 60-6583 [6]. These findings revealed that BAY 60-6583 administration significantly protected against liver injury in mice, even beyond the therapeutic window for NAC. Mechanistically, the study suggests that A2BAR activation preserves mitochondrial function despite c-Jun N-terminal kinase (JNK) activation and its translocation to the mitochondria. We demonstrated that APAP overdose triggers the translocation of A2BAR to the mitochondria. This translocation appears to be linked to JNK activation [7]. Notably, the study also suggests that A2BR interacts with the progesterone receptor membrane component 1 (PGRMC1) on the mitochondrial outer membrane, potentially influencing the activity of CYP2E1, the primary enzyme responsible for NAPQI formation. These findings highlight the potential of A2BAR agonists to influence upstream events after an APAP overdose and mitigate mitochondrial dysfunction and subsequent hepatocyte necrosis. However, from the clinical standpoint it is important to also assess liver recovery, since this can dictate patient prognosis, where robust liver regeneration facilitates survival after overdose and compromised recovery causes ALF.

Thus, the present study focuses on elucidating the mechanisms underlying the protective and regenerative effects of A2BAR activation by BAY 60-6583 during liver recovery after APAP-induced injury. By expanding on our previous work to include clinically relevant scenarios, we seek to identify novel downstream signaling pathways and cellular targets involved in A2BAR-mediated liver protection and repair, which could be leveraged for clinically relevant therapeutic approaches.

## 2. Material and methods

### 2.1 Animals and experimental design

Eight-week-old male C57BL/6J mice were purchased from Jackson Laboratories and housed in an environmentally controlled room with a 12 h light/dark cycle with free access to food and water at the University of Kansas Medical Center. All experimental protocols were approved by the Institutional Animal Care and Use Committee of the University of Kansas Medical Center and followed the criteria of the National Research Council for the care and use of laboratory animals. Before the experiments, mice were fasted overnight and administered APAP dissolved in warm saline at a dose of 300 mg/kg (Sigma-Aldrich, St. Louis, MO) intraperitoneally (i.p.), followed by BAY 60-6583. An initial dose response experiment determined that 4 mg/kg of BAY 60-6583 showed consistent effects (data not shown), and hence this dose was chosen for the study. Thus, mice were injected with 4 mg/kg BAY 60-6583 dissolved in 50% DMSO with PBS, or 50% DMSO with PBS alone as vehicle at 6 or 9 h after APAP treatment and euthanized under isoflurane anesthesia at 24, 48 or 72 h after APAP administration. Heparinized blood was collected and centrifuged at 18,000×*g* for 3 min to collect plasma. Liver was resected and either snap frozen in liquid nitrogen for subsequent storage at −80 °C, or fixed in 4% paraformaldehyde for histology.

### 2.2 Biochemical measurements and histology

Plasma alanine aminotransferase (ALT) activities were measured with the Point Scientific ALT Test kit (Point Scientific Inc., Canton, MI) as per the manufacturer’s instructions. Tissue for histology was fixed and embedded in paraffin. Hematoxylin and eosin (H&E) staining was performed for evaluation of the area of necrosis, which was quantified by comparison to the area of the whole section and expressed as a percentage.

### 2.3 Immunohistochemistry and immunofluorescence

Liver sections (5μm) were rehydrated and underwent heat-mediated antigen retrieval using a citrate buffer (pH 6.0). For immunohistochemistry, the sections were blocked in 3% BSA in serum followed by overnight incubation at 4°C with rabbit anti-mouse F4/80 antibody (Cell Signaling Technologies, #70076). Endogenous peroxidases were quenched by incubation in 3% hydrogen peroxide for 20 minutes. The tissue sections were then incubated in SignalStain Boost IHC Detection Reagent (HRP, Rabbit) (Cell Signaling Technologies, #8114) for 30 minutes. SignalStain DAB Substrate Kit (Cell Signaling Technologies #8059) was used as a detection reagent. For immunofluorescence, liver sections were incubated overnight with rabbit anti-mouse Perilipin-2 polyclonal antibody (Proteintech, #15294-1). The sections were then incubated with Alexa Fluor 488 Goat anti-rabbit IgG (ThermoFisher, A-11034) secondary antibody for 30 minutes. Imaging was done using a Nikon Eclipse Ti2 microscope.

### 2.4 Western blotting

Snap frozen liver tissue was homogenized in buffer (20 mM HEPES, pH 7.0; 2 mM EGTA, 1 mM EDTA, 1% Triton X-100, 10% glycerol, 150 mM NaCl, and 20 mM glycerol-2-phosphate) and centrifuged at 20,000×*g* at 4 °C for 5 min to remove cell debris and the supernatant separated. Protein concentration was measured using the BCA assay and samples were denatured at 95°C followed by separation on 10–12% SDS polyacrylamide gels and transfer to polyvinylidene fluoride (PVDF) membrane, which were then probed with primary antibodies at 1:1000 dilution overnight. Subsequently, the corresponding horseradish peroxidase (HRP) labelled secondary antibody (Cat. # sc2357 or sc2055; Santa Cruz Biotechnology, Dallas, TX) was used and proteins were visualized by enhanced chemiluminescence detection on a LICOR Odyssey Imager (LICOR Biosciences). The primary antibodies used were total Akt (pan) Rabbit mAb (#4691), Phospho-Akt (Ser473) Rabbit mAb (#4060), AMPKα Antibody (#2532), Phospho-AMPKα (Thr172) (40H9) Rabbit mAb (#2535S), GSK-3ß Rabbit mAb (#9315S), Phospho-GSK-3ß Rabbit mAb (#9323S) and GAPDH Rabbit mAb (#5174) from Cell Signaling Technologies (Danvers, MA).

### 2.5 Isolation of hepatocytes and non-parenchymal cells

Isolation of hepatocytes and non-parenchymal cells (NPCs) was performed at the COBRE Cell Isolation Core at KUMC using a protocol adapted from that of [9]. In brief, mice were anesthetized, and the liver was perfused via the portal vein with 25 ml of perfusion buffer (9.5 g/L Hanks’ balanced salt solution, 0.5 mmol/L EGTA, pH 7.2) followed by 50 ml of digestion buffer (9.5 g/L Hanks’ balanced salt solution, 0.14 g/L collagenase IV, and 40 mg/L trypsin inhibitor, pH 7.5).

After digestion, the livers were disaggregated by filtering through a 70 μm nylon mesh. The cell suspension was centrifuged at 50×*g* for 5 min to separate hepatocytes (pellet) from NPCs (supernatant). The supernatant was centrifuged at 350×*g* for 5 min. The cell pellet, now containing non-parenchymal cells, was then harvested and resuspended in 1 ml of red blood cell lysis solution (BD Biosciences). After incubation for 2 min, cells were washed once with 1% BSA buffer in PBS and centrifuged at 350×*g* for 5 min at 4 °C, following which the pellet was resuspended in BSA buffer for flow cytometry analysis.

### 2.6 Flow cytometric analysis of liver non-parenchymal cells

For flow cytometric analysis, hepatic non-parenchymal cells were first incubated with the Fcγ receptor blocker TruStain FcX™ antibody (anti-mouse CD16/32, Biolegend) to block non-specific binding of antibodies to Fc receptors. Cells were then stained with the following fluorescence-conjugated antibodies directed against the mouse, all purchased from Biolegend (San Diego, CA); CD45-PerCP (30-F11) (#103130), CD11b-PE-Cy5 (M1-70) (#101209), Ly6C-APC (HK1.4) (#128015), CX3CR1-PE (SA011F11) (#149006), Zombie Brillian Violet 510 (#423113) to identify live cells, and F4/80-FITC (#123108). The following marker combination was used for the definition of cell populations: Kupffer cells (KCs) were identified as CD45^+^ F4/80^hi^CD11b^lo^. Monocyte-derived macrophages were identified as CD45^+^ F4/80^lo^ CD11b^hi^. Anti-inflammatory monocyte-derived macrophages were identified as LyC6^lo^ CX3CR1^hi^ and the pro-inflammatory monocyte-derived macrophages were identified as LyC6^hi^ CX3CR1^lo^. The stained cells were analyzed on an LSR II flow cytometer (BD Biosciences) and the acquired data analyzed with FlowJo software (Tree Star).

### 2.7 RNA sequencing

Mouse hepatocytes and non-parenchymal cells were isolated as previously described from animals treated with APAP followed by BAY 60-6583 or 50% DMSO in PBS. RNA was extracted using TRIZOL reagent. RNA quantity and integrity were determined using the Agilent TapeStation 4200 using the RNA ScreenTape Assay kit (Agilent Technologies 5067–5576). Libraries were constructed utilizing the Universal plus mRNA-seq with NuQuant library preparation kit (Tecan Genomics 0520-A01) according to the manufacturer’s protocol. Paired-end sequencing was performed using the Illumina NovaSeq 6000 Sequencing System at the KUMC genomics core. Reads were aligned to the mm10 genome with HISAT2, indexed with Samtools, and a count matrix generated using Subread. Differentially expressed genes were determined using the DESeq2 package in R version 4.4.1. To investigate the biological functions of the differentially expressed genes (DEGs), Gene Ontology (GO) enrichment analysis was performed using the enrichGO function of the clusterProfiler R package (v4.0). Statistical significance was defined as an adjusted p-value < 0.05 [10].

### 2.8 Statistics

Data is expressed as means ± SEM. Comparisons between two groups were performed with Student’s *t*-test. Comparisons between multiple groups were performed using one-way ANOVA followed by Student-Neuman-Keuls post hoc test for multiple groups. The alpha level was set at

0.05 (P < 0.05).

## 3. Results

### 3.1 Activation of A2BAR by BAY 60-6583 promotes liver repair through Ras signaling pathway modulation

Our previous study indicated that activation of A2BAR by its agonist BAY 60-6583 protected against APAP overdose-induced liver injury by preventing mitochondrial dysfunction and, when administered at later stages, by increasing macrophage accumulation in the liver [6]. To further elucidate delayed mechanisms and effects of A2BAR activation by BAY 60-6583 following APAP overdose, we performed bulk RNA-seq analysis on liver samples collected 24 h post-APAP from animals treated with BAY 60-6583 at *6 h after* overdose. Initial analysis revealed the upregulation of 169 genes and downregulation of 303 genes in mice treated with APAP+BAY 60-6583 when compared with those treated with APAP + DMSO (vehicle control). Hierarchical clustering of the top 50 differentially expressed genes (DEGs) between the treatment groups was then visualized in a heatmap with dendrograms (Fig. 1A). Z-score normalized expression values were used to cluster both genes (rows) and samples (columns) based on expression similarity. The dendrogram revealed a clear separation between samples treated with APAP + DMSO and those treated with APAP + BAY, which indicates distinct treatment-specific transcriptional signatures. Genes upregulated or downregulated in the APAP+DMSO group (shown in red or blue respectively) showed an opposing regulation with BAY treatment, with APAP-upregulated genes being blunted and those suppressed by APAP being upregulated with BAY (Fig. 1A). Sample clustering also showed strong intra-group consistency, with all APAP + BAY samples forming a separate branch from APAP + DMSO samples, supporting a robust effect of BAY treatment on gene expression. Notably, several genes upregulated in the APAP + BAY group are involved in cAMP/GPCR and calcium signaling (e.g., *P2ry2, Gnao1, Cacng8, Pde4dip*) and in transcriptional regulation and cytoskeletal dynamics (e.g., *Zhx1, Trim54, Nucks1, Fubp3, R3hcc1l, Shoc2, Vti1a, Tc2n, Ralgapa2, Arhgap35*).

**Figure 1.**
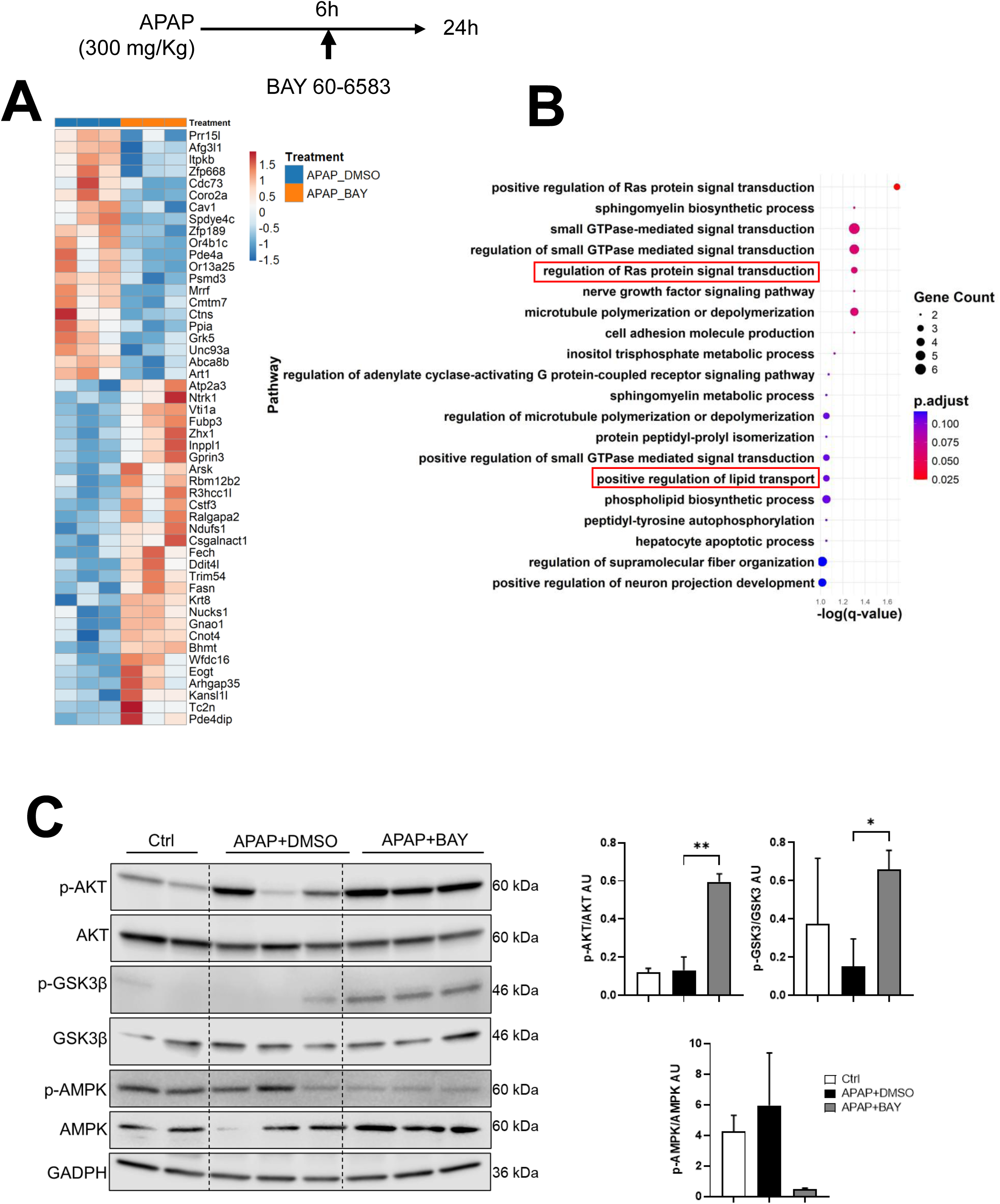
Transcriptional profiling reveals distinct gene expression signatures and enriched pathways between APAP+DMSO and APAP+BAY groups. Male mice were treated with 300 mg/kg APAP, followed by 4 mg/kg BAY 60-6583 or 50% DMSO (vehicle control) 6 h later. Bulk RNA sequencing from liver samples was performed 24 h post APAP overdose. (A) Hierarchical clustering heatmap of the top 50 differentially expressed genes (DEGs) between APAP+DMSO and APAP+BAY-treated livers at 24 h post-APAP overdose. Z-score normalized expression values are shown, with red indicating higher expression and blue indicating lower expression. Samples clustered distinctly by treatment group, suggesting treatment-specific transcriptional responses. (B) Pathway enrichment analysis of DEGs with log₂ fold change > 1. Top enriched pathways are shown, ranked by statistical significance (adjusted p-value). (C) Western blot analysis of Ras pathways and densitometry quantification. Data represent mean ± SEM, n = 3 animals per group. *p < 0.05 vs APAP + DMSO.

To further characterize the biological processes affected by BAY treatment, Gene Ontology (GO) enrichment analysis was performed on the DEGs. The analysis revealed significant enrichment of pathways related to intracellular signal transduction (Fig. 1B). Among the most significantly enriched terms was “positive regulation of Ras protein signal transduction” (q ≈ 0.025). Additional enriched terms included “sphingomyelin biosynthetic process” (q ≈ 0.03) and “small GTPase-mediated signal transduction” (q ≈ 0.032), the latter was notable for containing the largest number of contributing genes.

Collectively, the frequent enrichment of terms such as “small GTPase-mediated signal transduction”, “regulation of Ras protein signal transduction,” “microtubule polymerization or depolymerization,” “cell adhesion molecule production,” and “peptidyl-tyrosine autophosphorylation” highlights a consistent theme of transcriptional regulation affecting signaling and structural pathways. These results indicate that treatment with BAY 60-6583 leads to significant transcriptional modulation of genes involved in small GTPase signaling and cytoskeletal remodeling, strongly implicating these pathways in the compound’s protective effects and in the regulation of liver cell proliferation and repair post-APAP injury. We confirmed the activation of Ras protein signaling by evaluating the phosphorylation status of AKT, AMPK, and GSK3-β at 24 h after APAP overdose in animals treated with either APAP+BAY or APAP+DMSO (Fig. 1C). Animals treated with BAY 60-6583 exhibited significantly increased phosphorylation of both AKT and GSK3-β, and decreased phosphorylation of AMPKα compared to APAP+DMSO control group. This corroborates the pathway analysis findings and supports the role of BAY 60-6583 in activating Ras-mediated signaling cascades.

### 3.2 Effect of A2BAR activation on liver recovery post APAP overdose

Sterile inflammation plays a pivotal role in the response to liver injury and throughout the recovery process after a moderate APAP overdose. Although macrophages are essential for host defense, they can contribute to injury progression as well as tissue repair. Ly6C^high^ monocytes (pro-inflammatory) dominate within the first 24 h after APAP exposure but decline as regeneration progresses. Conversely, Ly6C^low^ monocytes (anti-inflammatory), initially present in low numbers, become the dominant population during the regenerative phase (48-72 h) [11, 12]. Our earlier data demonstrated that treatment with BAY 60-6583 not only protects against APAP-induced liver injury by reducing mitochondrial dysfunction but also modulates macrophage dynamics. Specifically, A2BAR activation enhances monocyte infiltration and helps to maintain the Kupffer cell population following APAP hepatotoxicity [6]. To understand how this shift in macrophage populations impacts liver recovery, we assessed liver injury and immune cell infiltration at 48 and 72 h post-APAP overdose in animals treated with BAY 60-6583 or DMSO 6 h after 300 mg/kg APAP administration.

Plasma ALT levels and necrotic area were evaluated at both time points. While ALT levels remained unchanged between groups (Fig. 2A), a significant reduction in necrotic area was observed in BAY-treated mice at both 48 and 72 h (Fig. 2B-C), reflecting enhanced recovery due to the protective effects of A2BAR activation. Immunostaining for F4/80 and flow cytometry analysis of immune response revealed that BAY 60-6583 treatment accelerated Kupffer cell replenishment at both time points (Fig. 3 A-B). Although the number of infiltrating monocytes did not differ significantly at 48 h post-APAP, a decrease was observed at 72 h (Fig. 3B) in the BAY 60-6583 treated group. The remaining infiltrating monocytes were predominantly anti-inflammatory or pro-regenerative monocytes (Ly6C^low^). These findings collectively suggest that A2BAR activation by BAY 60-6583 fosters a more rapid shift toward a pro-regenerative immune environment, supporting more effective liver recovery after APAP-induced injury.

**Figure 2.**
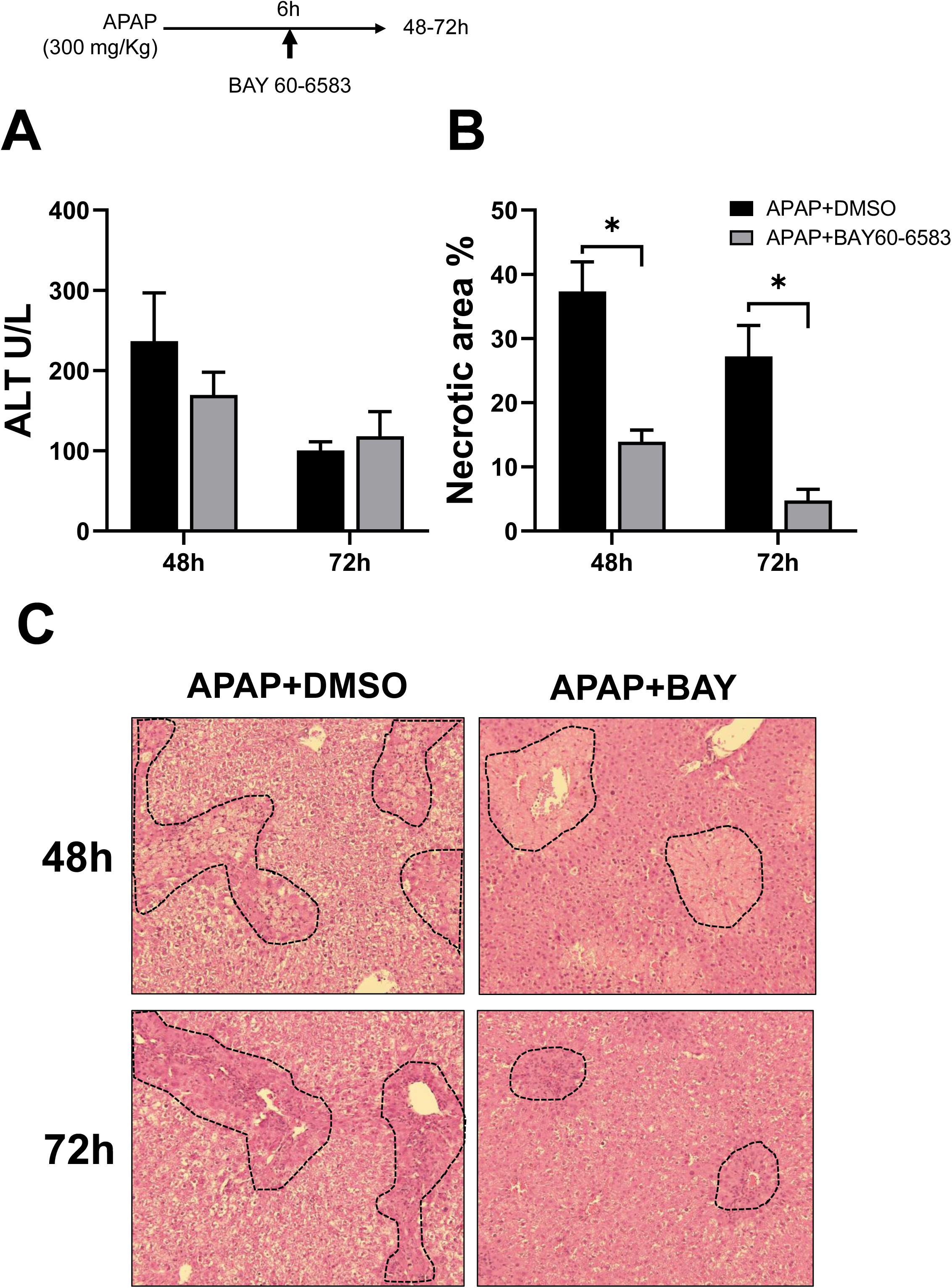
Delayed activation of A2BAR enhances liver recovery. Male mice were treated with 300 mg/kg APAP, followed by BAY 60-6583 6h later. Liver injury was evaluated at 48 and 72h after APAP overdose. (A) Plasma ALT, (B) Necrotic area quantification (C) H&E staining. Data represent mean ± SEM, n = 3-5 animals per group. *p < 0.05 vs APAP + DMSO.

**Figure 3.**
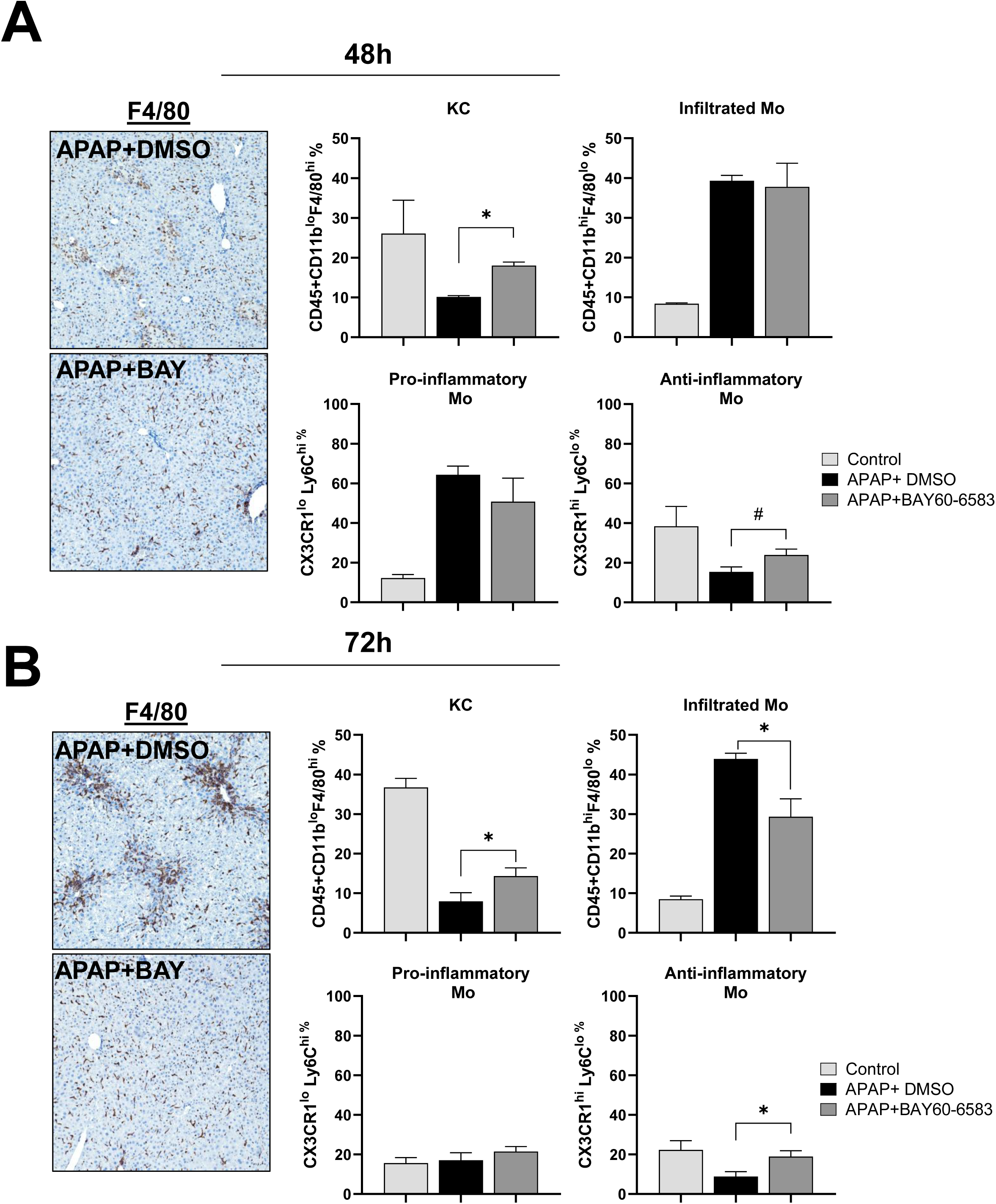
A2BAR activation by BAY 60-6583 promotes an anti-inflammatory immune response during liver recovery following APAP-induced liver injury. Male mice were treated with 300 mg/kg APAP, followed by 4 mg/kg BAY 60-6583 or 50% DMSO (vehicle control) 6 h later. Immunostaining of liver sections for F4/80 and flow cytometry analysis were performed at (A) 48 and (B) 72 h post APAP. Liver samples were obtained and flow cytometry analysis was performed at 48 (A) and 72 h (B) post APAP. Cell identification for flow cytometry was carried out as follows: Kupffer cells (F4/80^hi^, CD11b^lo^), infiltrating monocytes (F4/80^lo^, CD11b^hi^), pro-inflammatory monocytes (Ly6C^hi^, CX3CR1^lo^), and anti-inflammatory monocytes (Ly6C^lo^, CX3CR1^hi^). Bars represent mean ± SEM, n = 3-4 per group. *p < 0.05 vs APAP + DMSO.

Our data shows that treatment with BAY 6h post-APAP increased anti-inflammatory macrophage infiltration and helped to prevent KC depletion. To confirm that the pro-regenerative response and reduced areas of necrosis at 48 h were a direct effect of BAY 60-6583 treatment, and not due to any influence on the initial injury, we further delayed BAY 60-6583 treatment delayed to 9 h post-APAP administration, when injury processes should be complete. This also allowed us to further explore a broader therapeutic window for BAY 60-6583 (Fig. 4A). Animals treated with BAY at 9 h post-APAP showed no difference in plasma ALT levels or necrotic area at 24 h, indicating no effect on the initial phase of liver injury. However, by 48 h, while ALT levels remained unchanged, a reduction in necrotic area was observed in the BAY treated group (Fig. 4B & C), suggesting that A2BAR activation at this later time-point can still promote liver recovery. Evaluation of liver macrophages (Fig. 4D) showed that vehicle-treated animals exhibited depletion of Kupffer cells (KCs) by 24h, whereas BAY-treated mice maintained KC levels comparable to untreated control mice. A similar trend was also evident at 48h post APAP (Fig. 4D). Notably, there was no difference in the number of infiltrating monocytes at 48h, however there is an increase in anti-inflammatory monocytes, indicating a pro-resolution response. This suggests that BAY 60-6583 does not alter monocyte recruitment at 48h after a delayed treatment at 9 h post APAP but may specifically protect KCs from APAP-induced depletion. These findings highlight a role for A2BAR activation in preserving the liver’s resident macrophage population and facilitating tissue repair during the recovery phase of APAP-induced hepatotoxicity.

**Figure 4.**
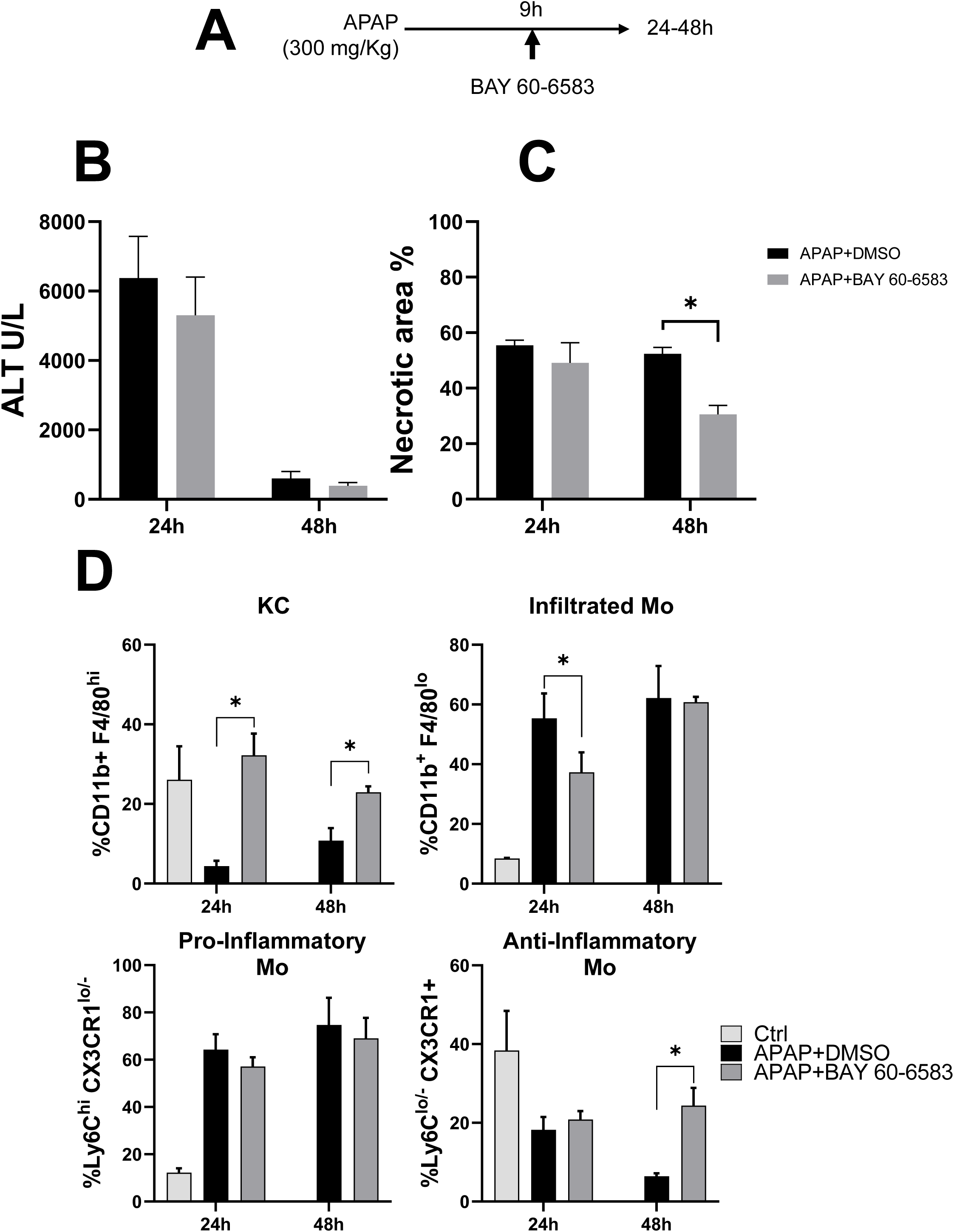
Very delayed activation of A2BAR at 9h post APAP also promotes a pro-resolution immune response. Male mice were treated with 300 mg/kg APAP, followed by 4 mg/kg BAY 60–658 or 50% DMSO as a vehicle control 9 h later. Samples were collected 24 and 48 h after APAP. (A) Experimental design. (B) ALT plasma levels. (C) Necrotic area quantification. (D) Flow cytometry analysis of hepatic non-parenchymal cells to evaluate Kupffer cells (F4/80^hi^ CD11b^lo^), infiltrating monocytes (F4/80^lo^ CD11b^hi^), pro-inflammatory monocytes (Ly6C^hi^ CX3CR1^lo^) and anti-inflammatory monocytes (Ly6C^lo^ CX3CR1^hi^). Bars represent means ± SEM, n = 3-5 per group. *p < 0.05 vs APAP+DMSO.

### 3.3 Distinct pathways activated by BAY 60-6583 in hepatocytes and non-parenchymal cells: proliferation and lipid metabolism in liver recovery

To further characterize the pro-regenerative environment generated by A2BAR activation and elucidate the regulatory gene networks in hepatocytes and non-parenchymal cell (NPC) during the recovery phase, we conducted bulk RNA-seq on both cell populations. RNA-seq data, collected 48 h post-APAP overdose, revealed distinct gene expression profiles between the APAP+DMSO and APAP+BAY 60-6583 groups in both NPCs and hepatocytes. Pathway analysis showed that NPCs exhibited upregulation of pathways associated with lipid metabolism, including lipid transport, regulation of lipid metabolic processes, and fatty acid degradation after BAY 60-6583 treatment (Fig. 5A-B). Conversely, hepatocytes exhibited increased expression of genes involved in cell proliferation, including pathways related to cell cycle transition, mitotic cell cycle, and DNA replication (Fig. 5C-D).

**Figure 5.**
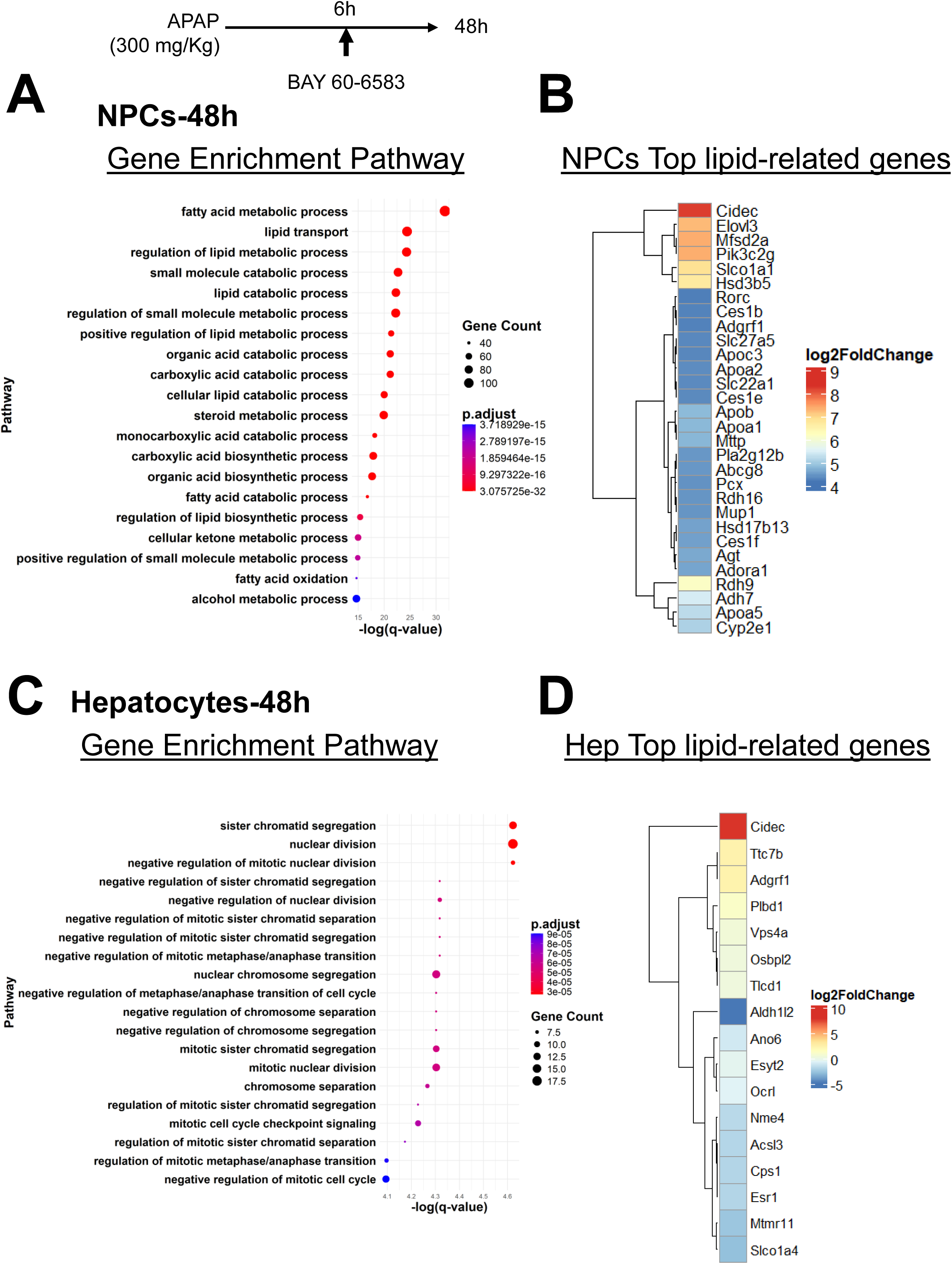
Transcriptomic analysis of hepatocytes and NPCs 48h post APAP in animals treated with 300 mg/Kg APAP followed by BAY 60-6583 6h later. (A) Gene enrichment pathways for NPC. (B) NPC-Top lipid metabolism related genes. (C) Gene enrichment pathway for Hepatocytes. (D) Hepatocytes-Top lipid metabolism related genes. Adjusted p value is obtained by comparing with APAP+ DMSO samples with n=3 for each group.

### 3.4 Regulation of lipid metabolism and the impact of BAY 60-6583 on liver recovery following APAP-induced injury

Through GO enrichment analysis, we identified that the most highly expressed pathway in NPCs was the fatty acid metabolic process (Fig. 5A). Following this analysis, we evaluated genes upregulated in lipid metabolism and found that *Cidec* was one of the most significantly upregulated genes in both NPCs and hepatocytes (Fig. 5B & D). *Cidec* (Cell Death-Inducing DFFA-Like Effector C) is a lipid transferase that plays a critical role in lipid metabolism, particularly in the regulation of lipid droplet (LD) formation and lipid storage [13]. Under normal physiological conditions, *Cidec* is not detectable in the liver; however, its expression increases during fasting or pathological states. Given the importance of *Cidec* in lipid droplet formation [14], we further evaluated other key genes associated with LD formation, such as *PLIN* genes [15]. Similar to *Cidec*, *PLIN2* and *PLIN5* were highly expressed in NPCs from animals treated with BAY 60-6583 (Fig.6A). Additionally, *PLIN2* and *PLIN4* expression were also significantly elevated in hepatocytes (Fig. 6B).

**Figure 6.**
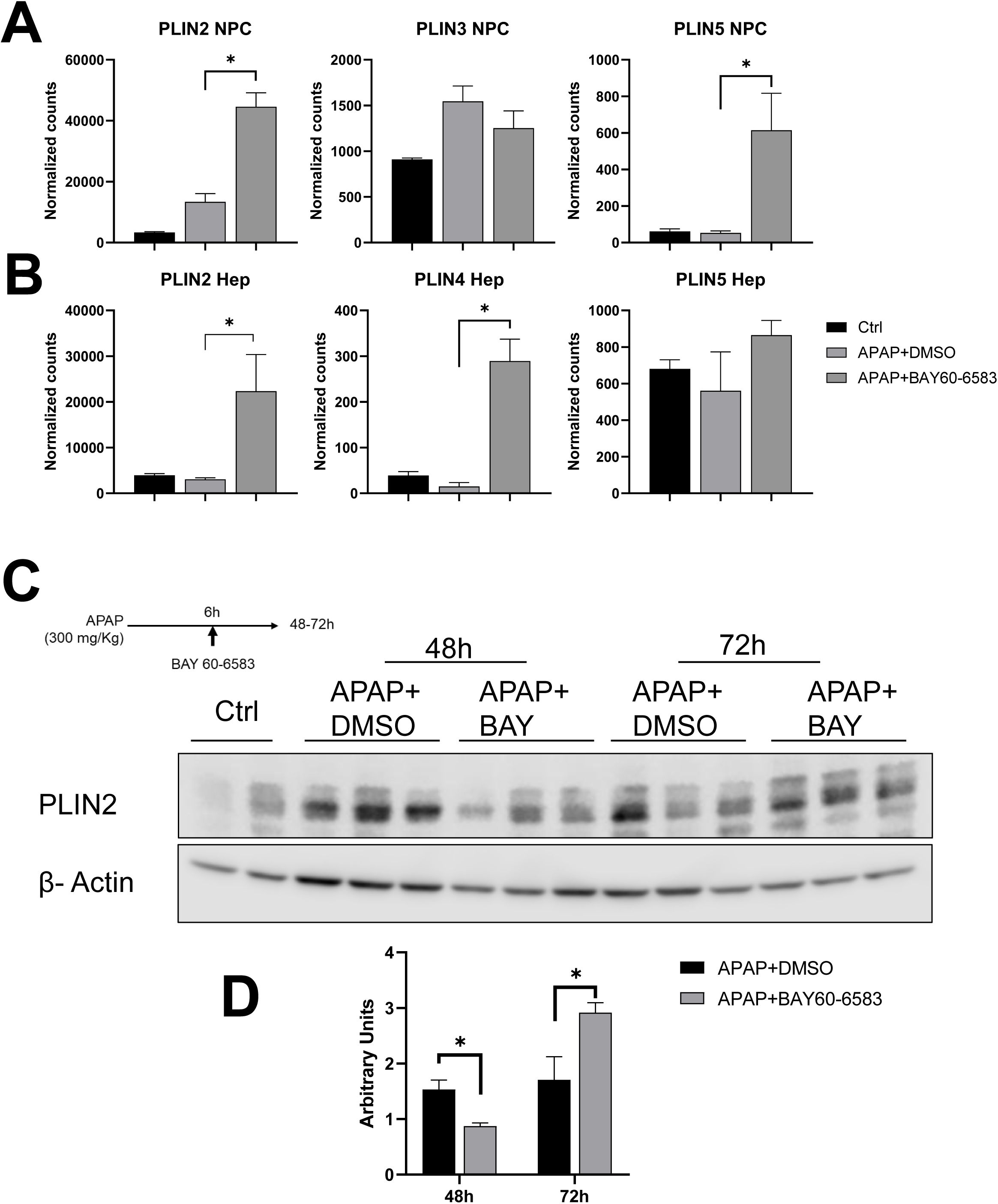
Perilipin- expression in hepatocytes and non-parenchymal cells. Perilipin gene expression using DESq2 normalized counts from the RNAseq data in non-parenchymal cells (A) and hepatocytes (B). (C) Western blot analysis of Perilipin-2 (PLIN2) protein levels in whole liver lysate. (D) Densitometry quantification of PLIN2 western blot. Bars represent means ± SEM, n = 3 per group. *p < 0.05 vs APAP+DMSO.

To corroborate these findings, *PLIN2* protein levels were analyzed via Western blot in whole liver lysates during the recovery phase after APAP overdose, with BAY 60-6583 administered at 6h post APAP. *PLIN2* protein levels were increased in the APAP+DMSO group at 48 and 72 h post-treatment compared to untreated animals (Fig. 6C-D). In contrast, animals treated with APAP+BAY 60-6583 showed significantly lower *PLIN2* levels at 48 h, with levels increasing by 72 h compared to the APAP+DMSO group (Fig. 6D). Lipid droplet formation in hepatic macrophages and hepatocytes is crucial for liver regeneration [14], as it contributes to free fatty acid (FFA) β-oxidation and energy supply. These results suggest that treatment with APAP+BAY 60-6583 enhances lipid metabolism, a finding that correlates with accelerated recovery following APAP overdose.

### 3.5 Perilipin-2 is expressed in perinecrotic hepatocytes

To investigate the spatial and cellular dynamics of PLIN2 expression during APAP-induced liver injury and recovery, we performed a series of transcriptomic and immunofluorescence analyses. We analyzed the expression of *Plin2* using the Visium spatial transcriptomics data from the Liver Cell Atlas (GSE280515) generated by Charlotte Scott’s group [16]. This dataset incorporates liver sections from mice treated with 300 mg/APAP and evaluated at 48 h post-APAP. *Plin2* mRNA expression was predominantly localized around the central vein area after APAP, suggesting a zonally restricted induction of PLIN2 expression following APAP exposure (Fig. 7A). To validate these spatial transcriptomic findings, we performed PLIN2 immunofluorescence staining on liver sections collected at 0 (Control), 24, and 48 h following APAP overdose (Fig. 7B). We observed minimal but homogeneous PLIN2 staining in control livers. At 24 h, PLIN2 expression markedly increased and was concentrated around the necrotic regions, particularly within perinecrotic hepatocytes. This zonal distribution persisted at 48 h, consistent with areas of ongoing injury and regeneration.

**Figure 7.**
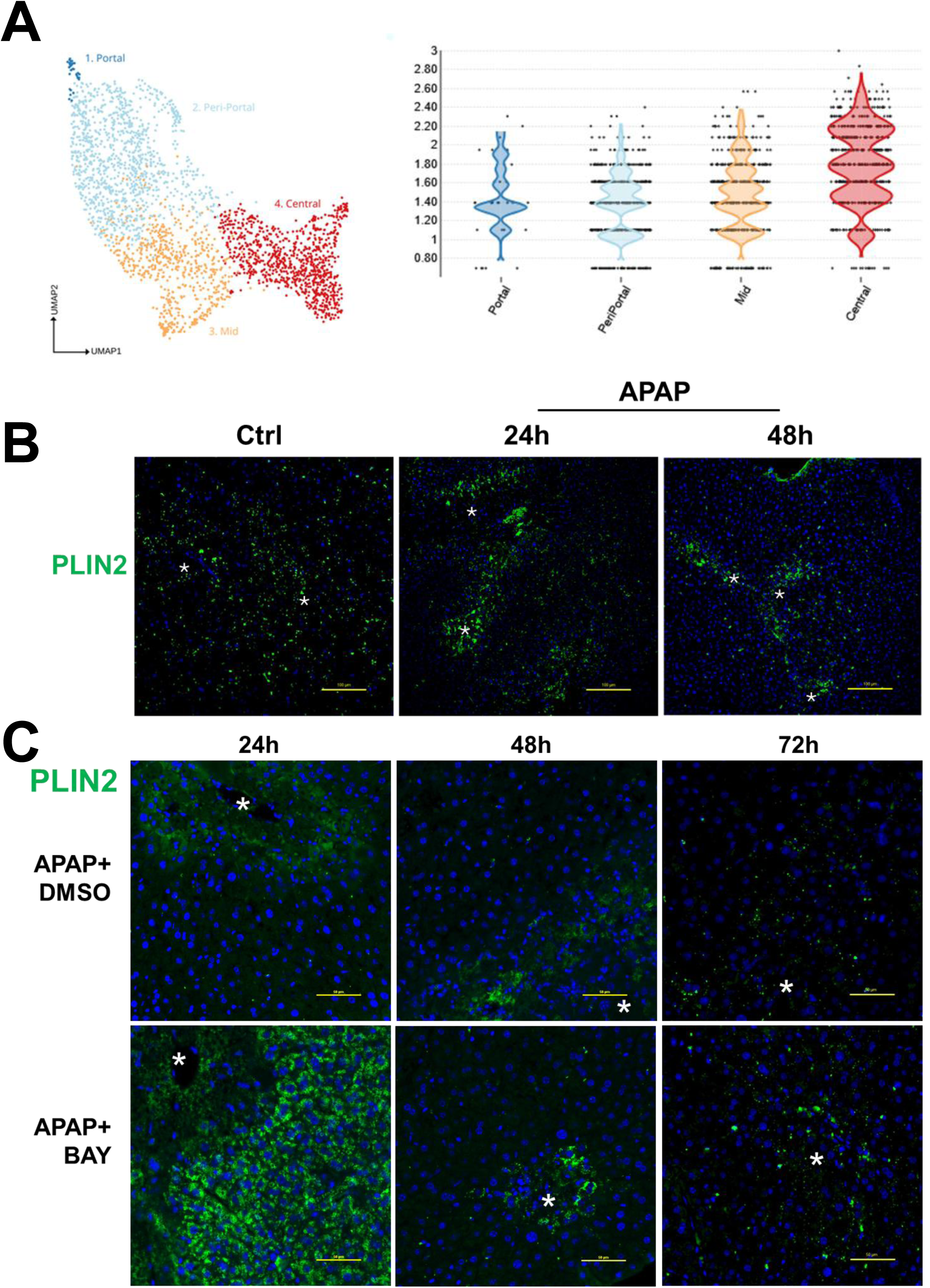
Activation of A2BAR induces perilipin-2 protein expression predominantly at 24h post APAP. (A) Plin2 Visium spatial transcriptomics analysis of livers from 48h APAP samples (https://www.livercellatlas.org). (B) Immunofluorescence staining of PLIN2 (green) in liver sections, expression evaluated at 24, 48, and 72h after APAP. (C) Immunofluorescence staining for PLIN2 in liver sections from mice treated with BAY 60-6583 at 6 h following APAP overdose. PLIN2 expression was evaluated at 24, 48, and 72 h post-APAP. * Marks the central vein.

To assess how A2BAR activation influences PLIN2 expression, we evaluated liver sections from mice treated with BAY 60-6583 at 6 h post-APAP. PLIN2 immunofluorescence staining was conducted at 24, 48, and 72 h post-APAP. At 24 h, we observed a widespread increase in PLIN2 expression across all liver zones in BAY-treated animals (Fig.7C). However, by 48 and 72 h, PLIN2 expression became more restricted to perinecrotic areas, and by 72 h, the expression pattern closely resembled that of untreated controls. These findings suggest that A2BAR activation modulates the temporal and spatial dynamics of PLIN2 expression, potentially contributing to the resolution of liver injury and the restoration of homeostasis.

## 4. Discussion

Despite significant advances in understanding APAP-induced hepatotoxicity, treatment options remain limited, highlighting an urgent need for novel therapies to enhance liver recovery and improve patient outcomes. Identifying mechanisms that can modulate sterile inflammation and facilitate regeneration following APAP overdose is therefore of critical therapeutic importance.

The beneficial role of the innate immune response after APAP-induced liver injury has been well documented [17–22], and involves three main mechanisms, including (i) clearance of necrotic cells from injured areas to initiate tissue repair; (ii) promotion of cellular proliferation through the production and release of various cytokines and growth factors; and (iii) production of pro-angiogenic factors thereby facilitating neo-vascularization [20]. It is well described that KCs are depleted 24h after APAP overdose [12], but the specific mechanism involved in KC depletion is still unknown. However, they are replenished during the recovery phase by proliferative KCs or bone marrow-derived monocytes (BMDM). Resident and infiltrated macrophages play an important role in the removal of the necrotic hepatocytes and promote liver regeneration. [11, 23, 24]. Several studies have highlighted the effects of A2BAR on regulating inflammation, especially on macrophage migration [25–27]. Our data are consistent with the hypothesis that this unique long-delayed protection by the activation of A2BAR through BAY 60–6583 treatment, even at 6 h or 9 h post APAP, is mediated through maintenance of Kupffer cell numbers and phenotype switching of infiltrating monocytes to the pro-regenerative phenotype. These changes in the innate immune response then facilitate liver regeneration as we had shown after netrin-1 treatment [8].

RNA sequencing revealed that hepatocytes treated with BAY 60-6583 showed upregulation of cell proliferation pathways at 24 and 48h post APAP. In contrast, NPCs exhibited enhanced expression of genes related to lipid metabolism, including fatty acid degradation and lipid transport. At 48 h post-APAP, PLIN2, a lipid droplet-associated protein [15], was highly expressed in perinecrotic hepatocytes, coinciding with the transition from injury to regeneration. This induction suggests an essential metabolic adaptation, providing energy and potentially modulating macrophage polarization towards a reparative phenotype.

A2BAR has previously been implicated in regulating lipolysis [28], glucose metabolism, and insulin sensitivity [25, 29, 30]. It has also been shown that A2BAR can regulate macrophage activation [25, 31], promotes anti-inflammatory macrophage polarization and induces IL-10 production [32–34].

Macrophage plasticity relies heavily on metabolic reprogramming and is context dependent. Pro-inflammatory macrophages rely on glycolysis associated with impaired oxidative phosphorylation (OXPHOS), whereas anti-inflammatory macrophages rely on the tricarboxylic acid (TCA) cycle, specifically enhanced fatty acid oxidation (β-oxidation) and enhanced mitochondrial oxidative metabolism [35–37]. Our data demonstrates a marked increase in lipid metabolism pathways in NPCs upon A2BAR activation, consistent with the emergence of anti-inflammatory macrophages that rely on fatty acid oxidation (FAO).

Recent studies have highlighted the critical role of distinct macrophage phenotypes during tissue repair, particularly emphasizing the importance of lipid-associated macrophages (LAMs) in promoting necrotic cell clearance and protection against APAP overdose [16, 38–40]. These LAMs are specifically enriched near necrotic areas, underscoring the significance of both macrophage phenotype and their zonal distribution for efficient tissue repair. Additionally, the LAM phenotype observed in Kupffer cells and infiltrating macrophages is transient and typically resolves once tissue regeneration is complete [16, 41].

The higher PLIN2 expression in the hepatocytes around the necrotic area, indicating lipid droplet formation, also correlates with liver repair and proliferation [42]. Transient hepatic steatosis, driven by elevated fatty acid uptake, is a hallmark of early liver regeneration, providing energy and anabolic substrates for proliferation. Disruption of this transient fat accumulation during early liver regeneration, such as genetic disruption of caveolin-1 and leptin, impairs regeneration following partial hepatectomy [43, 44]. Similarly, in the context of APAP hepatotoxicity, lipid metabolism also plays an important role in liver regeneration. For example, Mas activation in hepatocytes protects against APAP-induced liver injury by enhancing lipophagy and FAO [45].

Additionally, the inhibition of epidermal growth factor receptor (EGFR) impairs liver regeneration and eliminates lipid accumulation, suggesting that lipid metabolism is closely linked to the regenerative process and is regulated by EGFR signaling [46]. The absence of lipid accumulation after EGFR inhibition is attributed to the activation of AMPK-α, which inactivates acetyl CoA carboxylase (ACC), the rate-limiting enzyme of fatty acid synthesis, and decreases the expression of other enzymes involved in fatty acid synthesis [47]. Notably, A2BAR activation by BAY 60-6583 can induce EGFR phosphorylation and IL-6 production in skin inflammation models [9].

In our study, A2BAR activation decreased phosphorylation of AMPKα while increasing phosphorylation of ERK1/2, AKT, and GSK3β (Ser9), accompanied by enhanced PLIN2 gene and protein expression. These changes suggest a coordinated shift from catabolic to anabolic and lipogenic signaling. Reduced AMPKα activity suppresses fatty acid oxidation and autophagic lipid clearance [48, 49], while activation of ERK and AKT may promote the transcriptional activity of SREBP1 and C/EBPβ [50, 51]. Additionally, Inactivation of GSK3β by AKT-mediated phosphorylation further stabilizes β-catenin and SREBP1, reinforcing lipogenic gene expression, including PLIN2 [52–54].

This may explain the increased PLIN2 expression at 24 h post-APAP in BAY-treated animals, suggesting that transient lipid accumulation driven by A2BAR activation contributes to liver repair. However, further studies are necessary to elucidate the mechanisms by which A2BAR activation affects hepatocyte metabolism during regeneration.

In conclusion, in the context of APAP-induced liver injury, the activation of A2BAR with subsequent promotion of M2 macrophage polarization and enhanced lipid metabolism plays a critical role in facilitating liver regeneration. The metabolic reprogramming of macrophages toward FAO supports their anti-inflammatory functions, thereby aiding in the clearance of necrotic cells and the restoration of hepatic tissue. These insights underscore the therapeutic potential of targeting A2BAR signaling pathways to enhance liver repair mechanisms following toxic injury. Thus, A2BAR activation may be an adjunct therapeutic approach to NAC and fomepizole.

## Acknowledgements

This work was funded in part by the National Institute of Diabetes and Digestive and Kidney Diseases (NIDDK) grants R01 DK102142 (H.J.) and DK125465 (A.R.), and National Institute of General Medicine (NIGMS) funded Liver Disease COBRE grants P20 GM103549 (H.J.) and P30 GM118247 (H.J.). D.S.U. was supported by an NIH Predoctoral Fellowship (F31 DK134197). We thank the University of Kansas Medical Center Genomics Core for their support services, which is funded by the National Institutes of Health (NIH) funded Kansas Intellectual and Developmental Disabilities Research Center (NIH U54 HD 090216), the Molecular Regulation of Cell Development and Differentiation—COBRE (P30 GM122731-03), NIH S10 High-End Instrumentation grant (NIH S10OD021743), and the Frontiers CTSA grant (UL1TR002366) at the University of Kansas Medical Center.

